# The *Giardia lamblia* ribosome structure reveals divergence in translation and quality control pathways

**DOI:** 10.1101/2020.09.30.321331

**Authors:** Daniel R. Eiler, Brian T. Wimberly, Danielle Y. Bilodeau, Olivia S. Rissland, Jeffrey S. Kieft

## Abstract

*Giardia lamblia* is a human pathogen of worldwide importance with limited treatment options. Its unusual molecular biology presents targets for new therapies and the opportunity to explore the fundamental features of important biological mechanisms. We determined the structure of the *G. lamblia* 80S ribosome by cryoelectron microscopy, revealing how it combines eukaryotic and bacterial features. The structure reveals regions that are rapidly evolving, including depletion of A and U bases from its rRNA. Specific features of the *G. lamblia* ribosome suggest it is less prone to stall on problematic peptide sequences, and that the organism uses altered ribosome quality control pathways compared to other eukaryotes. Examination of translation initiation factor binding sites suggests these interactions are conserved despite a divergent initiation mechanism. This work defines key new questions regarding ribosome-centric biological pathways in *G. lamblia* and motivates new experiments to explore potential targetable mechanisms.

## INTRODUCTION

*Giardia lamblia* is a unicellular protist and the causative agent of giardiasis (Einarsson et al., 2016). Treatment of *G. lamblia* infection exploits the molecular differences between the pathogen and humans, but resistance to current therapies is increasing (Argüello-García et al., 2020). New therapeutic options are needed, which motivates further explorations into the unusual molecular machinery of *G. lamblia*. A deeply branching eukaryote of the Excavata supergroup (Figure 1A), *G. lamblia* uses the fundamental eukaryotic gene expression pathways, but its pathways and molecular machines are often simplified or streamlined in comparison with traditional model systems (Burki et al., 2020). Studies of these molecular differences will shed new insight into the biological and evolutionary diversity of eukaryotes.

**Figure 1.**
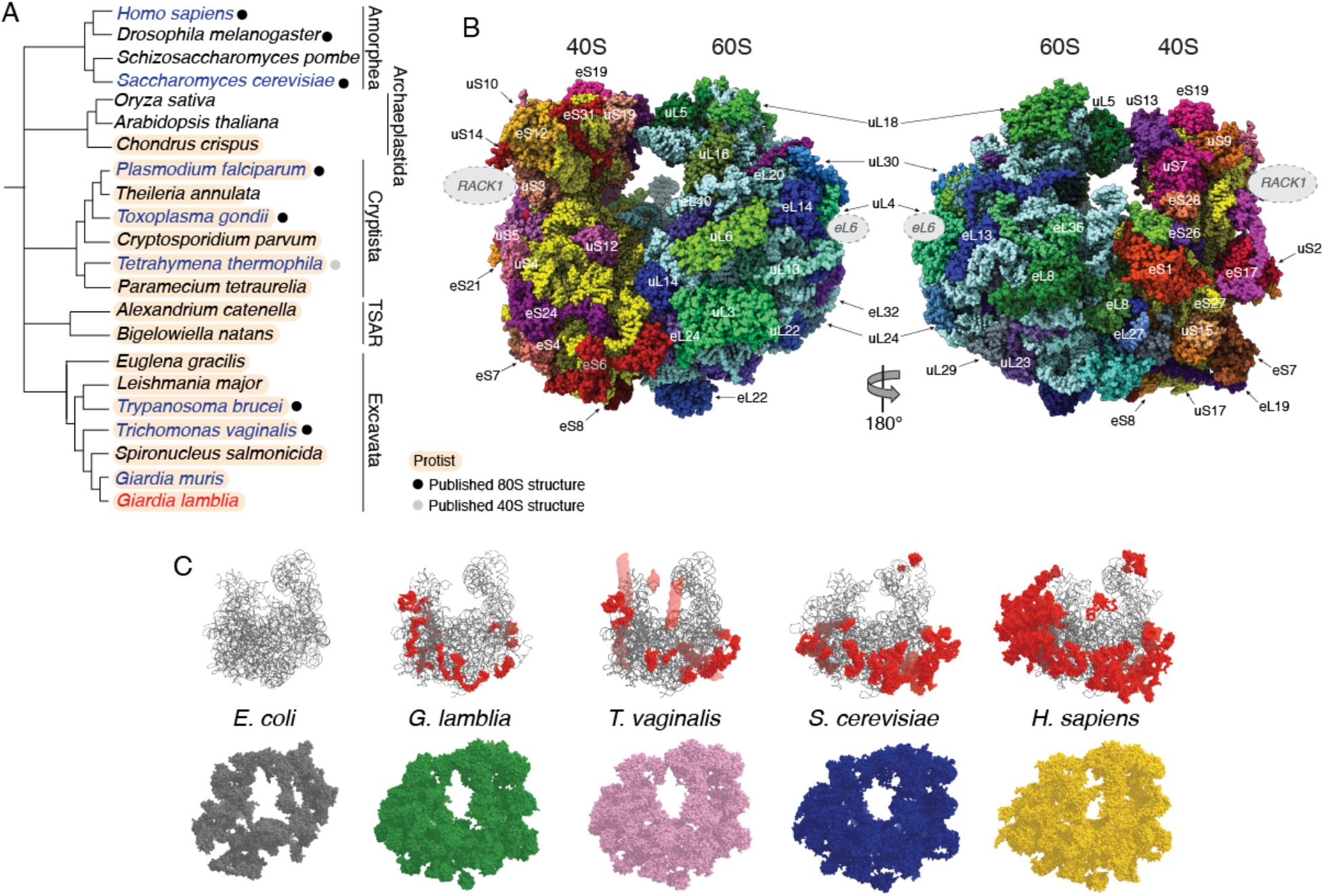
Architecture of the *G. lamblia* 80S ribosome. (A) Eukaryotic evolutionary tree. Shown is a schematized version of eukaryotic evolution with several of the protozoan branches. Species for which ribosome structures have been published are indicated with a dot. Protozoan species are highlighted in salmon. Names of species discussed here are in blue, except for *G. lamblia*, the focus of this study, which is in red. Branch lengths are not drawn to scale. (B) Two views of the *G. lamblia* 80S ribosome. 18S rRNA shown in gold and 40S ribosomal proteins (r-proteins) colored individually in generally ‘warm’ shades. 25S, 5.8S, 5S rRNA are in slight cyan and 60S r-proteins are colored individually in generally ‘cool’ shades. Proteins visible on the surface are labeled. The approximate positions of the missing ribosomal proteins RACK1 and eL6 are indicated with dashed grey circles. (C) Comparison of the rRNA and r-proteins between *E. coli* (PDB: 4U26), *G. lamblia, T. vaginalis* (PDB: 5XY3+5XYI), *S. cerevisiae* (PDB: 4V88), and *H. sapiens* (PDB: 5T2C). Top: The rRNA in each ribosome that is conserved across all species is shown as grey ribbons and eukaryotic rRNA expansion segments (ES) are shown as red spacefilling spheres. Transparent red surfaces on the *T. vaginalis* shows the estimated position and volume of rRNA ESs that were disordered in the solved structure. Bottom: The full set of r-proteins (without rRNA) for each ribosome is shown in a single color.

Translation, in particular, differs between *G. lamblia* and traditional eukaryotic model systems. In most eukaryotes, canonical translation initiation is cap-dependent and mediated through many eukaryotic initiation factors (eIFs) (Pelletier and Sonenberg, 2019; Shirokikh and Preiss, 2018). This includes the eIF4F complex, composed of the cap binding protein eIF4E, the large scaffold protein eIF4G, and the helicase eIF4A. The 5’ cap of the mRNA binds eIF4F though eIF4E, and interactions between eIF4G and the eIF3 complex recruit the 43S pre-initiation complex to the 5’ end of the message from where the ribosome begins scanning through the 5’ untranslated region (UTR). Upon start codon recognition, initiation factors are released, and the 60S subunit is recruited to form the 80S ribosome that commences elongation.

Although this process is stereotypical in almost all eukaryotes, nearly every aspect of translation initiation differs in *G. lamblia*—from the structure of its mRNAs, to a reduced cohort of initiation factors, to the ribosome itself. Like in other eukaryotes, mature *G. lamblia* mRNAs are capped and polyadenylated, but their UTRs are unusually short (Adam, 2000). For instance, most highly expressed transcripts have 5’UTRs shorter than 10 nucleotides, and reporters with a single nucleotide 5’UTR are readily translated in *Giardia* (L. Li and Wang, 2004). Moreover, lengthening 5’UTRs beyond nine nucleotides decreases translation, and functional studies suggest that *Giardia* ribosomes do not scan (L. Li and Wang, 2004). Thus, unlike in other eukaryotes, these results indicate a limited role for the 5’UTR in translation initiation in *Giardia*. Consistent with altered mRNA structure, *Giardia* also lack many eIFs almost universally conserved in eukaryotes and essential for life in yeast and humans (Ansell et al., 2019; Bannerman et al., 2018; Rezende et al., 2014). For example, there are no clear homologs for eIF4G and sequence analysis of *Giardia* eIF4E2 (its cap-binding protein) indicates that it does not have the consensus eIF4G-binding motif (L. Li and Wang, 2005), and recent studies suggest it can interact directly with other factors in translation preinitiation complexes (Adedoja et al., 2020). Together with an apparent lack of ribosome scanning, the absence of eIF4G points to a substantially reduced or even nonexistent role for eIF4F that may hint at other changes in the core translation machinery.

*Giardia* has also been reported to have major differences in the composition of the eIF3 complex, the largest initiation factor (Rezende et al., 2014). Based on sequence analysis, *Giardia* either lacks or has a highly divergent version of the eIF3a subunit, which is usually described as universally conserved (Gomes-Duarte et al., 2018; Valášek et al., 2017). In other organisms, the C-terminal tail of this large core subunit traverses the solvent side of the 40S ribosomal subunit and tethers the mobile eIF3b/i/g domain which does appear to be universally conserved. *Giardia* eIF3 also appears to lack subunits found in higher eukaryotes, including eIF3d, eIF3e, eIF3k, eIF3m, and eIF3l (Rezende et al., 2014), several of which have been found to bind eIF4A and/or eIF4G. These combined observations suggest a very different mode of ribosome recruitment and placement compared to other organisms and raise the question of whether there are corresponding changes in the ribosome.

As with other factors of the translation machinery, the *G. lamblia* ribosome differs from that in other eukaryotes. Its ribosomal RNA (rRNA) is over 3,000 nucleotides shorter than human rRNA and is the shortest of any known eukaryote except for a few microsporidia, and is even shorter than that of *E. coli* (Noeske et al., 2014). Nonetheless, *G. lamblia* ribosomes also have a nearly complete set of eukaryotic ribosomal proteins (r-proteins), and so its ribosome could be described as an amalgam of the bacterial and eukaryotic ones that is likely to have both similarities and differences compared to previously solved 70S and 80S ribosome structures (Anger et al., 2013; Ansell et al., 2019; Ben-Shem et al., 2011; Selmer et al., 2006). Because the ribosome acts as a central hub in post-transcriptional gene regulation, differences in ribosome structure could affect processes that recognize “aberrant” ribosome structures, such as the no-go decay (NGD) surveillance pathway (Simms et al., 2017). No-go decay is triggered by collided ribosomes (called “disomes”), which form when a trailing ribosome runs into a leading ribosome that has encountered an elongation block. Surveillance machinery then recognizes the interface between the two collided ribosomes, triggering destruction of the transcript and nascent peptides. Notably, a recent report indicates that another protist, *Plasmodium falciparum*, fails to recognize classic no-go decay substrates, raising the question of whether other protists might also lack this surveillance pathway (Pavlovic-Djuranovic et al., 2020). However, with the ribosome structure from *G. lamblia* and many other protozoa species unsolved (Figure 1A), these possibilities remain unexplored.

Motivated by these long-standing mysteries, we used cryo-electron microscopy to solve the structure of the *G. lamblia* 80S ribosome with the goal of providing a framework for future investigations. We obtained maps to an overall resolution of 4.02 Å, allowing nearly the complete ribosome structure to be built and examined. Differences in the structure compared to those of other eukaryotic ribosomes highlight areas of apparent rapid evolutionary change in both rRNA and proteins, compared to ribosomes of even closely related species. For example, in addition to being smaller, the rRNA is also depleted of adenines and uridines and appears to have only maintained these nucleotides when necessary for a specific structural role. Changes in both surface and interior features of the *G. lamblia* ribosome hint at changes in ribosome biogenesis pathways and interactions with nascent peptides. Surprisingly, these ribosomes also lack both rRNA and protein features associated with the specific inter-ribosome interface important for triggering no-go decay, raising the possibility that this surveillance pathway has been lost in at least two protists. Overall, this structure provides foundation and motivation for ongoing studies to understand the molecular biology of an evolutionary distant and important human pathogen.

## RESULTS AND DISCUSSION

### Global features of the *G. lamblia* 80S ribosome: a mix of bacterial and eukaryotic features

We determined the structure of the *G. lamblia* 80S ribosome at an overall resolution of 4.02 Å using single-particle cryo-electron microscopy (cryo-EM) with data obtained exclusively from a 200 keV microscope and direct electron detector (Figure 1–figure supplement 1). In addition to the rRNA, we observed density corresponding to most expected proteins based on the annotated genome (Figure 1B). We also observed clear density for protein eL41, which is only 24-25 amino acids in length and presumably due to its small size, is not annotated in any of the ten available *G. lamblia* sequenced genomes (Figure 1–figure supplement 2). We therefore used the sequence of yeast eL41 in our structural model.

Although most r-proteins were observed, we were unable to identify density for either protein RACK1 or eL6, which are both conserved proteins (Figure 1B). To validate these observations, we performed mass spectrometry analysis on two separate preparations of purified *G. lamblia* 80S ribosomes (Figure 1–Source Data 1, 2). The results confirmed the presence of all r-proteins except eS19, eS30, eL39, eL40, and eL41, but density for these were readily visible in the map at locations matching those in yeast and *T. vaginalis* 80S structures. In contrast, the mass spectrometry did not detect eL6 and RACK1, consistent with a lack of any corresponding features in the map. Together these results reveal that despite the small size of its rRNA, the *G. lamblia 80S* ribosome contains the full complement of eukaryotic r-proteins except for eL6 and RACK1.

The other notable difference in the *Giardia* ribosome is the absence of many rRNA expansion segments (ESs), which, while expected from the rRNA sequence, is striking in the structure (Figure 1C). While human ribosomes have 30 named ESs with many located on the periphery of the 80S structure, *Giardia* have only 13 ESs totaling 356 nt, similar in number and size to those in *T. vaginalis* (Anger et al., 2013; Z. Li et al., 2017). Consistent with its reduced rRNA and missing ESs, the *G. lamblia* 80S ribosome is also smaller in overall size compared to other eukaryotic ribosomes. The smaller *G. lamblia* rRNA is also reflected in its set of intersubunit bridges, which are more like bacteria than other eukaryotes (Figure 1–figure supplement 2). Thus, overall the *G. lamblia* ribosome melds bacteria-like and eukaryotic-like features.

### *G. lamblia* rRNA is dramatically depleted of adenines and uridines

Although much of the *G. lamblia* rRNA matches that of ribosome structures from other species, some portions differed, even when compared to other protists in the Excavata group such as *T. vaginalis*. First, in 25S rRNA, the region corresponding to nucleotides 448-498 has expanded compared to *T. vaginalis* (Figure 2A). Although not as large as it is in yeast or human (which have ES9L), the *G. lamblia* insertion has structural features similar to these (Anger et al., 2013; Ben-Shem et al., 2011; Z. Li et al., 2017). Second, a more striking difference is at the 3’ end of 25S rRNA, corresponding to nucleotides 2559-2606 (Figure 2B). In *G. lamblia*, this region is ~50% cytidines with many protein contacts and base stacking, but it has very few RNA-RNA hydrogen bonds or Watson-Crick base pairs. The result is a smaller more ‘contorted’ fold in *G. lamblia* compared to that in yeast, human, and *T. vaginalis*, although the filled space and overall shape of the region is similar to that of yeast and human. The 3’ end of the 25S rRNA also varies across *Giardia* species: for example, it is much shorter in *G. muris* (Figure 2–figure supplement 1). These characteristics suggest that this region is a rapidly evolving element on the solvent accessible surface of the large subunit rRNA.

**Figure 2.**
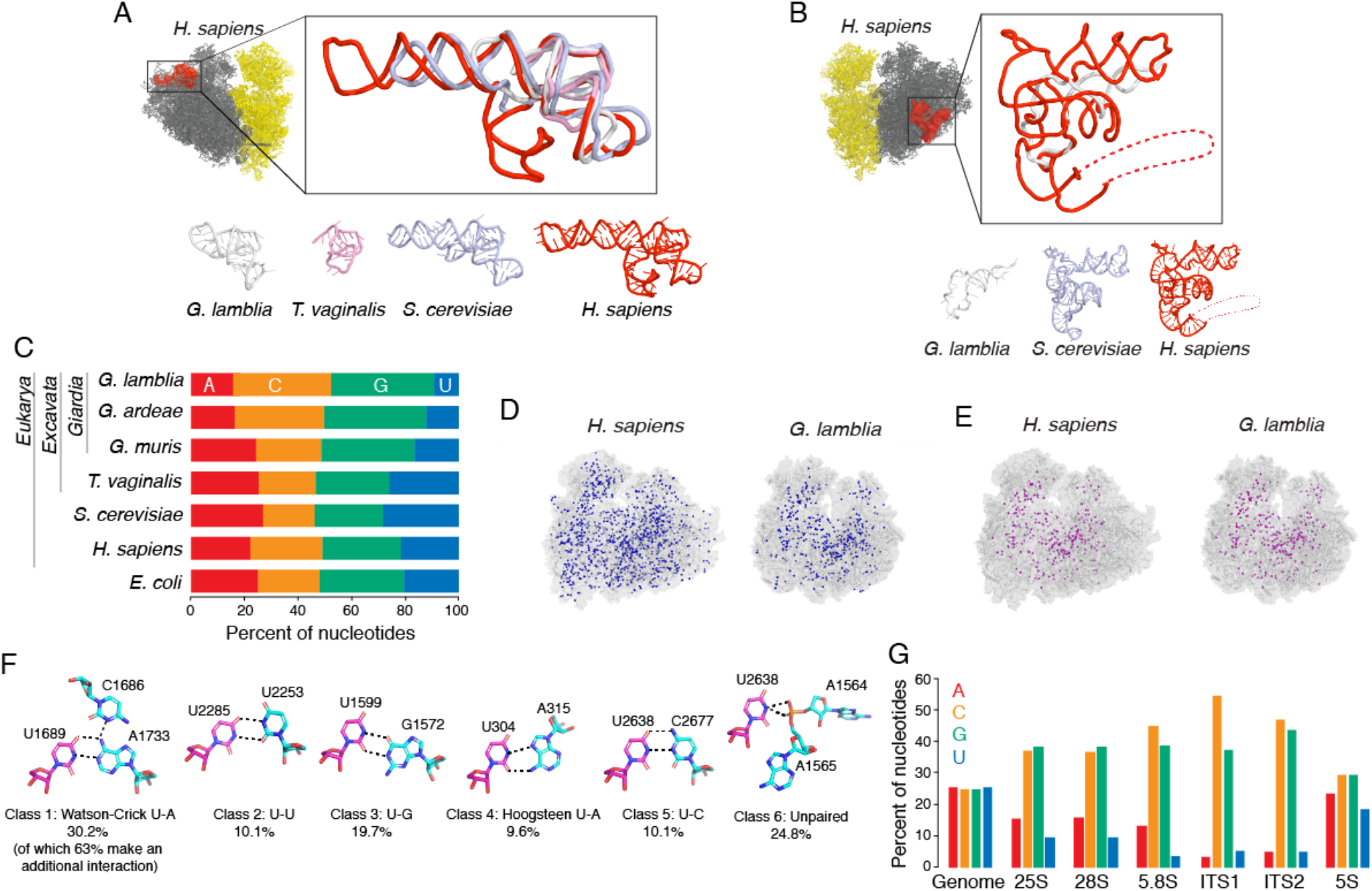
Features of the *G. lamblia* rRNA that contrast with other species. (A) Differences in the 28S rRNA between *G. lamblia* and other species. Left: *H. sapiens* 80S ribosome (60S in grey, 40S in yellow) with a region of the 28S rRNA that includes ES9L in red and boxed. Boxed: Overlay of this region of rRNA from *G. lamblia* (light gray), *T. vaginalis* (pink), *S. cerevisiae* (pale blue), and *H. sapiens* (red). Below: This region of rRNA from each species displayed side by side. (B) Differences in the 28S rRNA between *G. lamblia* and other species. Differences Left: *H. sapiens* 80S ribosome (60S in grey, 40S in yellow) with a region of the 28S rRNA that includes ES39L in red and boxed. As in A, except this region of rRNA in *T. vaginalis* is larger than in *G. lamblia* but is almost completely disordered, precluding structural comparison. (C) Comparison of the nucleotide composition of rRNA between *G. lamblia* and other species. Shown is the fraction of each nucleotide in mature rRNA. (D) Comparison of uridines between the *G. lamblia* and *H. sapiens* ribosomes. All rRNA uridines are indicated by blue spheres in *H. sapiens* (left) *and G. lamblia* (right). (E) Comparison of conserved uridines between the *G. lamblia* and *H. sapiens* ribosomes. The subset of uridine positions that are conserved between *H. sapiens* (left) and *G. lamblia* (right) are shown as magenta spheres. (F) Structural classifications of uridines in the *G. lamblia* ribosome. Examples of the interactions made by uridines in the *G. lamblia* ribosome with the percentage of uridines found in each class included. (G) *G. lamblia* polymerase I transcripts are depleted in adenines and uridines. Comparison of the nucleotide percentage in different RNAs produced in *G. lamblia*. 25S, 18S, 5.8S are transcribed by RNA polymerase I with the ITS sequences being part of immature rRNA and removed during rRNA processing. 5S RNA is transcribed by RNA Polymerase III.

The marked nucleotide bias towards cytidine in the 25S rRNA 3’ end (50%) prompted us to explore the nucleotide composition of the *G. lamblia* rRNA as a whole. The G/C content of the entire *G. lamblia* genome is 46%, but its rRNA is 75.3% (Figure 2C). Although human rRNAs are also G/C rich (65.2%), much of this nucleotide bias resides in ESs, while the G/C content is only 54.1% in the central ‘core’ of the human rRNA shared with *G. lamblia*. The strikingly high G/C content of the *G. lamblia* rRNA means that the compositions of A and U are only 15.3% and 9.4%, respectively. This dramatic depletion of uridines is noteworthy as to our knowledge, no other ribosome has <10% of any one nucleotide. This skewed nucleotide composition is not seen even in *G. muris* (Van Keulen et al., 1993) and only to a lesser extent in *G. ardeae* (Figure 2C), suggesting a recent and rapid shift towards A/U-poor rRNA in *G. lamblia*.

To investigate this nucleotide bias further, we focused on uridine loss because it showed the strongest signal. Our structure of the *G. lamblia* 80S ribosome allowed us to determine if this depletion of uridines was associated with specific parts of the structure. In fact, within the rRNA, the distribution of remaining uridines is uniform, indicating a reduction in the use of this base throughout the rRNA but with specific positions being maintained (Figure 2D, E). These maintained positions are mostly conserved compared to the human ribosome, but 60% of the uridines found in human rRNA were a different nucleotide in *G. lamblia*, and most often cytidines (Figure 2–figure supplement 2). We next considered two non-exclusive possibilities why these Us are maintained. First, some U bases might be important for post-transcriptional modification (e.g., pseudouridylation). Because modification sites have not been mapped in *Giardia* rRNA we were unable to assess this directly, so we used the sites of modification in human ribosomes as a guide and examined the structurally analogous locations in the *G. lamblia* rRNA. This analysis revealed no compelling evidence that potential sites of modification are preferentially maintained as uridine (Figure 2–figure supplement 2). A second possibility is that some uracil bases might make important non-Watson-Crick tertiary interactions that cannot be formed by other bases. Consistent with this possibility, examination of the structure showed that relatively few uridines in the *G. lamblia* rRNA are found in Watson-Crick base-pairs that do not also make additional uracil-specific structural interactions (Figure 2F). We saw similar trends when we looked at adenine residues, though adenines are conserved mostly in non-Watson-Crick pairing or other tertiary interactions. While the moderate resolution of our structure precludes making a conclusion for every uridine and adenine, it appears that those not required for a specific structural role by the nucleobase have largely been replaced.

We next investigated why *G. lamblia* has replaced its A and U rRNA bases, considering two possible reasons. There could have been a selection to replace A–U pairs with G–C base pairs to stabilize the overall rRNA structure; alternatively, there may be some unidentified transcriptional limitation (*e.g*., the availability of nucleotides for making new ribosomes). To tease apart these possibilities, we examined the nucleotide composition in the ITS1 and ITS2 spacer regions that are transcribed as part of the unprocessed rRNA precursor (K. E. Bohnsack and M. T. Bohnsack, 2019; Henras et al., 2008). If the nucleotide shift in mature rRNA is due to pressure to stabilize the rRNA structure specifically, then the spacer regions should roughly match the nucleotide composition of the overall genome rather than that of the mature rRNA. Contrary to this prediction, however, the ITS1 and ITS2 spacer regions were even more depleted of A/U nucleotides, with a G/C percentage of 89.4% (Figure 2G). In other words, because the spacer regions have even fewer structural constraints than mature rRNA, they presumably have freedom to move towards even higher G/C enrichment. Thus, the G/C enrichment is unlikely to be explained by a ribosome structure-based evolutionary pressure, but rather by some broader transcriptional effect. Interestingly, the bias against A/U seems limited to only RNA Polymerase I as non-coding transcripts from other polymerases did not show this same effect (Figure 2G). Although the underlying mechanism is unknown, the fact that rapidly evolving regions such as the 3’ end of 25S and the ITS regions are particularly G/C-rich, and that this bias is not seen in the closely related *G. muris* species suggests the presence of an unidentified evolutionary pressure on RNA polymerase I leading to a nucleotide bias in *G. lamblia*.

### Important eIF3 binding sites are conserved

Previous analyses suggested that *G. lamblia* lacks several eIFs including some subunits of the large and dynamic eIF3 complex. In humans and other higher eukaryotes, eIF3 contains an ‘octamer core’ (subunits a, c, e, f, h, k, l, and m), peripheral subunits (d, j, and n), and a ‘yeastlike core’ (YLC) (subunits b, i, g, and the C-terminal tail of a); in contrast, many lower eukaryotes such as *S. cerevisiae* have only a subset of these (YLC, j, a and c) (Gomes-Duarte et al., 2018; Valášek et al., 2017). In *G. lamblia*, candidates for subunits b, c, f, g, h, i, and j have been identified but not the a subunit (Rezende et al., 2014), which is surprising given its nearly universal conservation and the role of its C-terminal tail in tethering the universally conserved and highly mobile YLC 3b/i/g domain.

We used our structure to explore whether the altered G. lamblia eIF3 was reflected in its potential interactions with the 40S subunit. To this end, we used the recent structure of a human 48S preinitiation complex to construct an exploratory model (Brito Querido et al., 2020). To accomplish this, we superimposed the *G. lamblia* 40S subunit with the 40S subunit in the 48S structure, and then threaded candidate individual eIF3 subunit sequences through the corresponding structures (Figure 3A, B). As subunits b, c, and d are among the most important for anchoring the eIF3 complex to the 40S subunit, we examined the corresponding modeled interaction surfaces. In general, the *G. lamblia* 40S subunit is very similar to human in these regions, although there are some changes in the size of rRNA ESs near the subunit binding sites. Specifically, in *G. lamblia* ES7S is missing between subunits a and c, while ES6S is missing and h16 is truncated adjacent to subunit b (Figure 3B). Because *G. lamblia* has no identified eIF3a, we looked to see if the corresponding binding surface on the 40S subunit has changed from humans. In fact, the surface is well-conserved structurally (Figure 3A, B). This result, together with the observation that *G. lamblia* appears to have several eIF3 subunits that interact with 3a in other organisms, suggests that there is an 3a subunit in *G. lamblia* whose divergent sequence precludes easy identification. The identification of additional ‘missing’ *G. lamblia* eIF3 components, the verification of current candidates, and the implications of changes in the presence or size of ESs on factor binding are important foci of future explorations.

**Figure 3.**
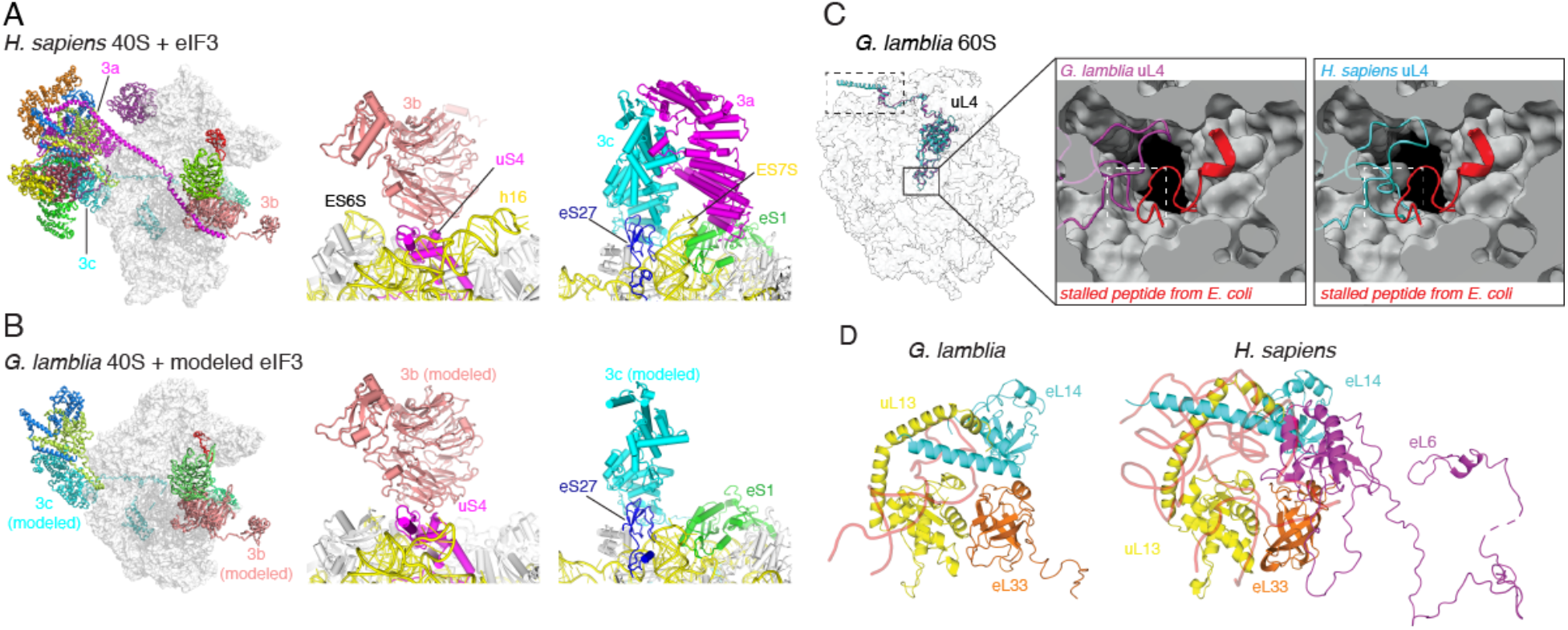
Comparison of *G. lamblia* r-proteins and ribosome interaction surfaces with other species. (A) Structure of the human 40 subunit (transparent surface) with eIF3 bound (various colors), constructed from PDB entry 6ZMW (Brito Querido et al., 2020). Important eIF3 subunits that contact the 40S are labeled. Middle: Close-up view of the interaction between eIF3b and the 40S subunit, with subunit features labeled. Right: Close-up view of interactions between eIF3a, eIF3c and the 40S subunit. (B) Modeled interactions between the *G. lamblia* ribosome and eIF3. Left: modeled structure of the *G. lamblia* 40S subunit with the eIF3 subunits proposed to exist (Rezende et al., 2014; Xu et al., 2020). Subunit models were generated using the Phyre2 server (Kelley et al., 2015), with sequences from the GiardiaDB database (https://giardiadb.org/giardiadb/app) (see Methods). Middle: Modeled interactions between eIF3b and the 40S subunit. Right: Modeled interactions between eIF3c and the 40S subunit. Note that a candidate for eIF3a has not been identified in *Giardia*, but the binding surface of the subunit on the 40S subunit matches that in humans (panel A, right). (C) The *G. lamblia* ribosome has a less constricted exit channel. Left: Structure of the *G. lamblia* 60S subunit is shown as a transparent surface with uL4 from *G. lamblia* (purple). and human (cyan). Dashed box: location of an N-terminal truncation in *G. lamblia* relative to human. Solid box: location of a loop truncation in *G. lamblia* in the 60S subunit’s peptide exit tunnel. Right: The peptide (red) from a structure of a stalled *E. coli* ribosome (PDB entry 6TC3; (Herrero Del Valle et al., 2020)), modeled in position to show the path of the nascent peptide until it reaches uL4. *G. lamblia* uL4 is in purple, human uL4 is in cyan. The white dashed box indicates the location of a loop that differs between the two species. (D) Altered substructure of uL13, eL14, and eL33 in the *G. lamblia* ribosome. Left: Interactions between uL13 (yellow), eL14 (cyan), eL33 (orange), and rRNA (transparent red) in the *G. lamblia* 60S subunit. Right: This same region in *H. sapiens*, in which eL6 (purple) is present. eL6 inserts one domain between eL33 and eL14, inducing a shift in the position of parts of eL14 which propagates to uL13. Human ribosome is PDB entry 5T2C (Zhang et al., 2016).

### Features on the interior and exterior of *G. lamblia* ribosomal proteins diverge from other eukaryotes

Our structure also revealed changes both on the exterior and in the interior of the *G. lamblia* ribosome. One example is protein uL4, which lines part of the peptide exit tunnel and extends out to the solvent side of the large subunit (Figure 3C). Within the interior of the subunit, *G. lamblia* uL4 has a loop that is 6 amino acids shorter than does *H. sapiens*, yeast, and *T. vaginalis* uL4. This loop is located in a constricted point in the tunnel where specific nascent peptide sequences have been shown to stall and halt ribosome elongation (Figure 3C) (Han et al., 2014; Herrero Del Valle et al., 2020). The uL4 truncation in *G. lamblia* therefore may change the characteristics of this part of the exit tunnel, potentially reducing the probability that the ribosome stalls on ‘problematic’ peptide sequences. Interestingly, *G. lamblia* uL4 is also truncated by 31 amino acids on its C-terminus. In the Tetrahymena, *H. sapiens* and yeast ribosome structures, this part of uL4 forms an alpha helix that wraps around the solvent side of the large subunit, but in *G. lamblia* this region is missing (Figure 3–figure supplement 1). Because uL4 is involved in ribosome biogenesis (Fox et al., 2019; Lawrence et al., 2016), this result may indicate changes in the formation of *G. lamblia* ribosomes.

In addition to uL4, other proteins on the surface of the *G. lamblia* ribosome differ from what has been observed in other ribosome structures. One is caused by the absence of eL6, which is present in all previously-solved eukaryotic ribosome structures (Z. Li et al., 2017). The structural effects of lacking eL6 can be seen by comparing this region of the *G. lamblia* and *H. sapiens* structures. In humans, eL6 is buried by rRNA ES39L and to a lesser degree by ES7L, but both are missing from *G. lamblia* leading to structural changes involving eL14, uL13, and eL33 (Figure 3D). In humans, uL13 and eL14 interact through their C-termini, and eL33 does not interact with eL14 because a part of eL6 is wedged between them (Figure 3D). In *G. lamblia*, however, the uL13 and eL14 C-termini shift such that eL14 contacts eL33, and uL13 fills in the volume where eL14 is in humans and other eukaryotes. The implications of these changes in this ‘substructure’ of the *G. lamblia* 80S ribosome are unknown, but as eL14 has been implicated in ribosome biogenesis, this result adds additional weight to a hypothesis that this process is altered in *G. lamblia* (Feng et al., 2020).

### *G. lamblia* 80S ribosome structure suggests altered quality control pathways

A notable feature of the *G. lamblia* 80S ribosome is the absence of protein RACK1, although it was easily identifiable in the 80S structure from the closely related protist *T. vaginalis*. This observation could stem from several causes: (1) RACK1 could have dissociated during ribosome purification, (2) *G. lamblia* RACK1 might not bind to ribosomes, or (3) *G. lamblia* may not even express RACK1. To explore these possibilities, we searched the *G. lamblia* genome for RACK1 candidates using both the structural homology search tool Phyre2 and the protein sequence BLAST tool with RACK1 sequences and structures from other eukaryotes (Kelley et al., 2015; Z. Li et al., 2017). We identified two candidate *G. lamblia* proteins with structural homology to yeast RACK1 — however, RACK1 contains a beta-propeller type fold found in many proteins and thus the best candidate was not clear (Figure 4—figure supplement 1). However, based on RNA-seq analyses, both candidates were expressed at substantially lower levels compared to other ribosomal proteins (Figure 4A). Thus, while we cannot unambiguously determine whether *G. lamblia* has a RACK1 homolog, we suggest the lack of a RACK1-like protein may be an authentic feature of the native ribosome in the trophozoite stage.

**Figure 4.**
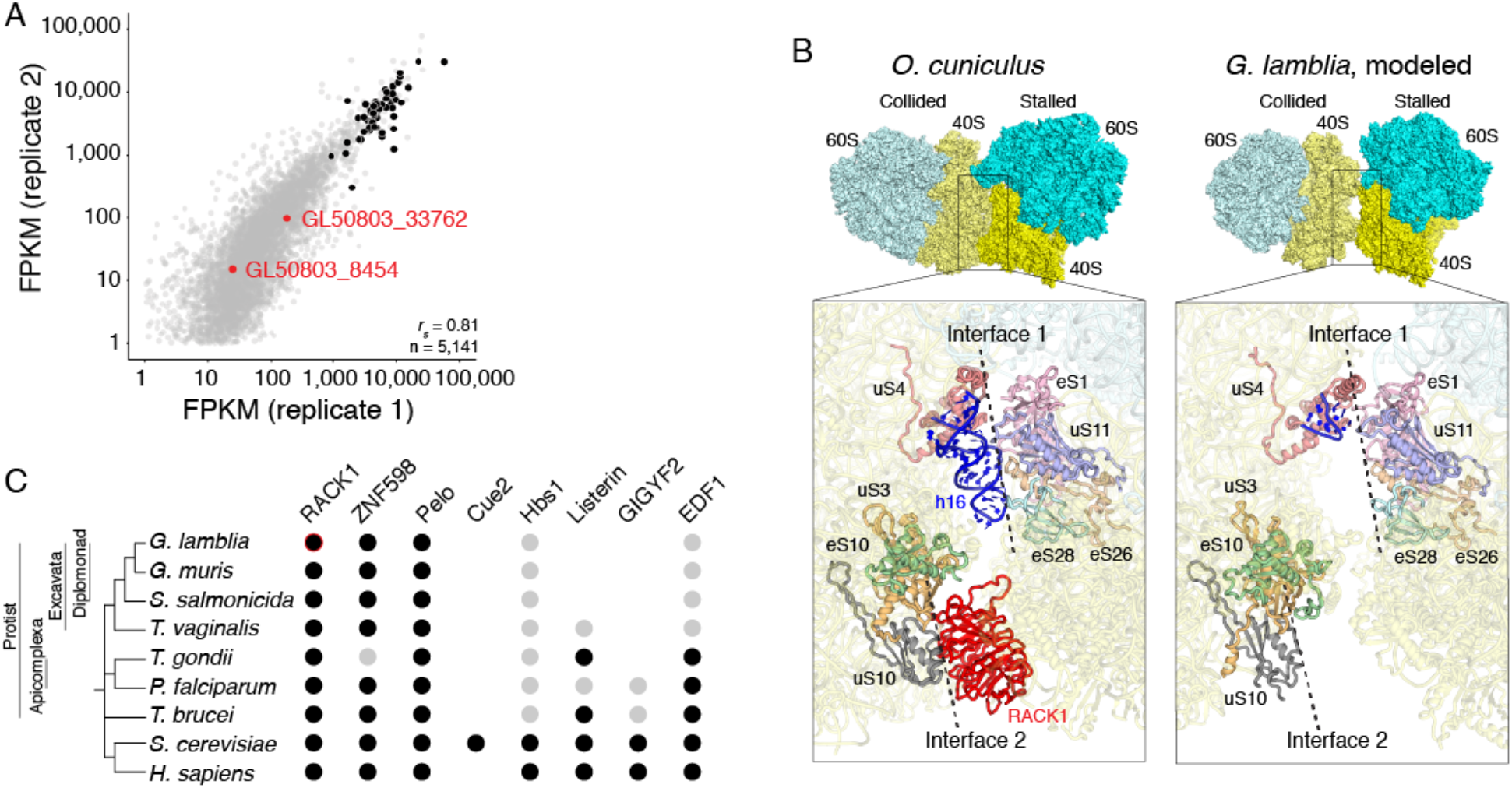
*G. lamblia* 80S structure reveals potential differences in quality control pathways. (A) Potential RACK1 transcripts are lowly expressed. Shown is a scatter plot from independent RNA-seq experiments. Transcripts encoding r-proteins are shown in black; the two potential RACK1 orthologs are shown in blue. (B) Comparison of authentic disomes from rabbit (left) and modeled *G. lamblia* (right). Top: surface representation with 40S shown in shades of yellow and 60S in shades of cyan. Bottom: close-up of the interaction surface, highlighting proteins previously identified as participating in the inter-ribosome interactions and the two named interfaces (Juszkiewicz et al., 2018). RACK1 and h16 are present in the rabbit 80S and form interactions within the rabbit disome, but these features are lacking in the modeled *G. lamblia* disome. (C) Identification of putative orthologs of ribosome quality-control pathway factors. Potential orthologs of RQC factors were identified in various protists using BLAST. Orthologs for which there was strong hit are marked with a black dot; orthologs with weak evidence are shown with a grey dot. RACK1 is highlighted in red due to evidence that it does not associate with *G. lamblia* ribosomes in the trophozoite stage.

Interestingly, 80S ribosome structures from two other protists (*Plasmodium faliciparum* and *T. gondii*) have also lacked RACK1 (Z. Li et al., 2017; Wong et al., 2014). Transcriptomics data available suggest that the *P. falciparum* ortholog of RACK1 is highly expressed in multiple life stages, leading to the idea that a *bona fide* RACK1 is expressed, but its association is weak enough to be lost during sample preparation. Nonetheless, these observations prompted us to examine the interface between RACK1 and the rest of the ribosome. In many structures, the C-terminal tail of uS3 appears to help anchor RACK1 to the 40S subunit. The length of this tail varies between species, but those with the shortest tails (such as *T. gondii*) have lacked RACK1 in the resulting structures (Figure 4—figure supplements 2, 3). Interestingly, *G. lamblia* uS3 has the shortest C-terminal tail known to date. This tail thus has shortened several times in different protozoa lineages, which raises the possibility that the strength of the association of RACK1 with the ribosome has changed substantially throughout eukarya.

RACK1 is linked to many physiological processes in humans such as tumor growth, apoptosis, and neural development (Duff and Long, 2017; Kershner and Welshhans, 2017; Nielsen et al., 2017), and at the molecular level is crucial for ribosome quality control pathways, such as no-go decay (NGD). Here, RACK1 was recently shown to be part of the interaction between two collided ribosomes within ‘disomes’ that trigger ribosome-dependent quality control pathways (Ikeuchi et al., 2019; Juszkiewicz et al., 2018). Prompted by the lack of RACK1, we constructed a model of what the *G. lamblia* disome might look like. By docking the *G. lamblia* 80S structure into both copies of the 80S ribosome in a rabbit disome structure (Juszkiewicz et al., 2018), we observed predicted changes to the interface between the two ribosomes (Figure 4B). Strikingly, a number of previously identified key interactions are lost in both inter-ribosome interfaces including interactions involving RACK1 and h16 (Ikeuchi et al., 2019; Juszkiewicz et al., 2018), the latter of which is truncated in *G. lamblia*. Thus, colliding ribosomes in *G. lamblia* likely cannot form disome structures in the same way as do their human hosts; disomes, if they form, must use a different interface and rely on different inter-80 ribosome contacts.

Interestingly, when we examined the *G. lamblia* genome for proteins involved in NGD and RQC, we were unable to identify orthologs for several key proteins, including Listerin and GIGYF2, in *G. lamblia* and other protists (Figure 4C). Given the recent report that classic NGD substrates fail to elicit decay in *P. falciparum* (Pavlovic-Djuranovic et al., 2020), our results raise the idea that this pathway is surprisingly plastic throughout eukaryotic evolution and the implications warrant further investigation.

### Conclusions and outlook

The structural model of the *G. lamblia* 80S ribosome presented here provides the first view of the basic translation machinery of this evolutionary-distant human pathogen, revealing differences that provide the basis for new hypotheses. The structure supports the fact that several key molecular biology pathways are altered in this organism, including ribosome biogenesis, translation initiation, and ribosome quality-control pathways. In addition, changes in the nucleotide composition of the rRNA indicate potentially significant and evolutionary recent changes in the function of *G. lamblia* RNA polymerase I, suggesting an unknown but potentially strong selective pressure. Additional biochemical, structural, and genetic explorations spurred by these discoveries promise important insights to guide future developments of new therapies.

## MATERIALS AND METHODS

### Trophozoite culture and collection

*G. lamblia* trophozoites (assemblage A, strain WB Clone C6) were purchased from the ATCC and grown as per standard protocols. To scale up growth for collection of ribosomes, trophozoites were grown in 10-cm culture dishes and incubated at 37°C for 48 hours within a GasPak EZ Container system with fresh anaerobic sachets. To harvest the cells, media was removed from the plates and cells were scraped and collected in 1XPBS, then spun down for 5 minutes at 1,500 x g at 4°C. The cell pellet was washed once more in 1XPBS and stored at −80°C.

### RNA sequencing and analysis

RNA was extracted from trophozoites with hot acid phenol as previously described (Collart and Oliviero, 2001). Poly(A)-selected RNA was used to generate libraries with Illumina TruSeq stranded mRNA library preparation kit as per the manufacturer’s protocol. Libraries were sequenced at The Centre for Applied Genomics at The Hospital for Sick Children in Toronto, Canada. Reads were mapped to the *G. lamblia* genome (release-43, 2019-04-19) using STAR 2.5.2a and gene abundance was quantified using Cufflinks 2.2.1 (Dobin et al., 2013; Trapnell et al., 2010). Downstream analysis was completed with RStudio from in-house scripts. Sequencing data are available from the GEO (accession number GSE158187).

### Purification of 80S ribosomes

Frozen cell pellets of *G. lamblia* were suspended in lysis buffer (150 mM KCl, 15 mM MgOAc, 15 mM Tris-HCl pH 7.4, 0.5% NP40, 1 mM DTT, 1 μl RNAsin, Roche protease inhibitors without EDTA). The cells were lysed by one freeze-thaw cycle in large volume (15–50 ml) and sheared by forcing 4X though a 25-gauge needle. Cell debris was cleared by centrifuging at 16,000 x g for 15 min at 4°C. Supernatant was layered over a 30% sucrose cushion (7.5 ml) in buffer A (20 mM Tris-HCl pH 7.4, 2 mM MgOAc, 150 mM KCl) and ultracentrifuged tor 16 hr at 36K (50.2 Ti Rotor) to pellet the ribosomes. The pellet was suspended in buffer A and ultracentrifuged through a 15–30% sucrose gradient in buffer A at 25K for 11 hr (SW28 rotor). The gradient was fractionated and monitored for absorption at 260 nm. Fractions containing the 80S ribosome peak were collected, then buffer A was added to dilute the sucrose and the sample was ultracentrifuged for 16 hr at 36K (50.2Ti) to pellet the ribosomes. The pellet was washed to remove any traces of sucrose and then resuspended in buffer A with 6 mM MgOAc, 1 mM DTT, and 1 μl RNAsin Plus.

### Cryo-EM sample preparation and data acquisition

Purified *G. lamblia* 80S ribosomes were used at a concentration of 15 μM. Using a FEI VItrobot, a 3 μl aliquot was applied to plasma-treated holey carbon grids (C-flat Cu 400 mesh) covered with graphene oxide. Grids were incubated for ~3 s before blotting for 2.5–3.5 s, then plunged into liquid ethane. Grids were stored in ice-free liquid nitrogen at least overnight before use. Grids were then transferred to a Thermo Scientific Talos Arctica microscope operated at 200 kV and equipped with a Gatan K3 direct electron detector. 8119 movies of 20 frames were collected in counting mode at 51.58 e^-^/pix/s at a magnification of 36,000X corresponding to a calibrated pixel size of 1.4 Å. Defocus values specified in Leginon ranged from 0.5 to 5.0 μM (Carragher et al., 2000). Data collection was monitored and checked during collection using APPION (Lander et al., 2009).

### Image processing and structure determination

Image processing was carried out in parallel using both Relion and CryoSPARC (Punjani et al., 2017; Scheres, 2012). Particles were picked at first manually using a particle diameter of 350-400 Å and used as a training set in automatic picking. Particles were then placed into 2D Classification using 40–200 classes per run to filter out lone subunits, ice, aggregates, and any other debris. All good classes were placed into *ab initio* 3D classification searching for up to 5 classes at a time. Initial reconstructions at lower resolution (7-9 Å) revealed features about the position of L1 stalk and an E-site-bound tRNA. Refinement improved the overall map, however the part of the maps containing the L1 stalk and the bound tRNA became featureless. Attempts to use hetero-refinement of the particles into classes that could separate out the positions of the L1 stalk and presence or absence of the Esite tRNA were not successful. These techniques only teased out different rotated states of the small ribosomal subunit relative to the large ribosomal subunit. Focused refinement of the 40S ribosomal subunit specifically improved the map in the head region. Final refinement of the map for the 60S and body of the 40S was done in Relion. Final refinement of the map for the 40S head was done in CryoSPARC.

### Model building and refinement

The initial model for the *G. lamblia* 80S ribosome was generated by using i-Tasser to generate models for the ribosomal proteins using *T. vaginalis* and yeast 80S ribosome structures as templates. Due to the lack of a single complete sequence source, the 25S rRNA was built using sequence information from GenBank: X52949.1 and Giardia DB: DHA2_r061, GL50803_r0021, GL50803_r0013, and GL50803_r0016 combined with interpretation of the density maps. Ribosomal RNA for the model was generated using Assemble2 and the *Trichomonas vaginalis* rRNA as a template. These initial models aligned to the *T. vaginalis* ribosome subunit structures and then rebuilt in COOT to using the Relion and CryoSPARC maps (Emsley et al., 2010). rRNA chains were refined using ERRASER after initial placement and adjustment (Chou et al., 2013). Refinement of the individual ribosomal subunits was done in Phenix (Liebschner et al., 2019).

### Modeling Giardia lamblia 40S subunit interactions with eIF3

Candidate genes for eIF3 subunits in *G. lamblia* were taken from (Rezende et al., 2014) and (Xu et al., 2019). 3b: GL50803_15495; 3c: GL50803_24279; 3f: GL50803_7896; 3g: GL50803_13269; 3h: GL50803_16823; 3i: GL50803_13661; 3j: GL50803_15546. To generate the model shown in Figure 3B, our structure of the *G. lamblia* 40S subunit was docked into the structure of the 48S complex from (Brito Querido et al., 2020), PDB entry 6ZMW using the ‘align’ command in Pymol to superimpose the 18S rRNA. Models of individual *G. lamblia* eIF3 subunits generated using the Phyre2 server (Kelley et al., 2015). Briefly, individual sequences downloads from the GiardiaDB database (https://giardiadb.org/giardiadb/app) were threaded into individual subunits from PDB entry 6ZMW using the online ‘One-to-one threading’ tool in Phyre2 ‘Expert Mode.’ Resultant coordinate files were combined with the 40S subunit structure without further modification to generate the model and images of Figure 3B.

### Ortholog identification

Human protein sequences were used to search for orthologs in the species of interest by BLAST search. Where it was difficult to identify the most likely ortholog among the search results, the yeast protein sequence was used for a complimentary search. For Listerin, GIGYF2, and EDF1, protist orthologs that were more readily identified (ex. *Trypanosoma brucei* or *Toxoplasma gondii)* were also used to BLAST against Trichomonas, Giardia and Spironucleus genomes to help identify the most likely ortholog. Searches were conducted on NCBI’s protein BLAST tool and various VEuPathDB websites: giardiadb.org, trichdb.org, toxodb.org, tritrypdb.org and plasmodb.org.

## DATA DEPOSITION

The coordinates for the 60S and 40S ribosomal subunits of the *G. lamblia* 80S ribosome were deposited in the protein data bank with the accession codes 7K9F and 7K9G, respectively. RNA-seq data are available from the Gene Expression Omnibus, accession number GSE158187.

## ACKNOWLEDGEMENTS

We thank Peter Van Blerkom (University of Colorado Anschutz Medical Campus Cryo-EM Facility) for assistance with microscope operation, and members of the Asturias Lab and Dr. Erik Hartwick for advice with data analysis and structure calculations. We thank members of the Kieft and Rissland labs for incisive conversations, and Dr. Aaron Jex and Dr. Staffan Svärd for assistance with *G. lamblia*. This work was supported by NIH grants R35GM118070 (JSK), R21AI149210 (OR and BW), and R35GM128680 (OR).

## SUPPLEMENTARY MATERIAL

**Figure 1–Source Data 1. Mass Spectrometry analysis of purified *G. lamblia* 80S ribosomes.**

Supplied as a separate spreadsheet file.

**Figure 1–Source Data 2. *G. lamblia* r-protein reference table.**

Supplied as a separate spreadsheet file.

**Figure 1–Figure Supplement 1.**
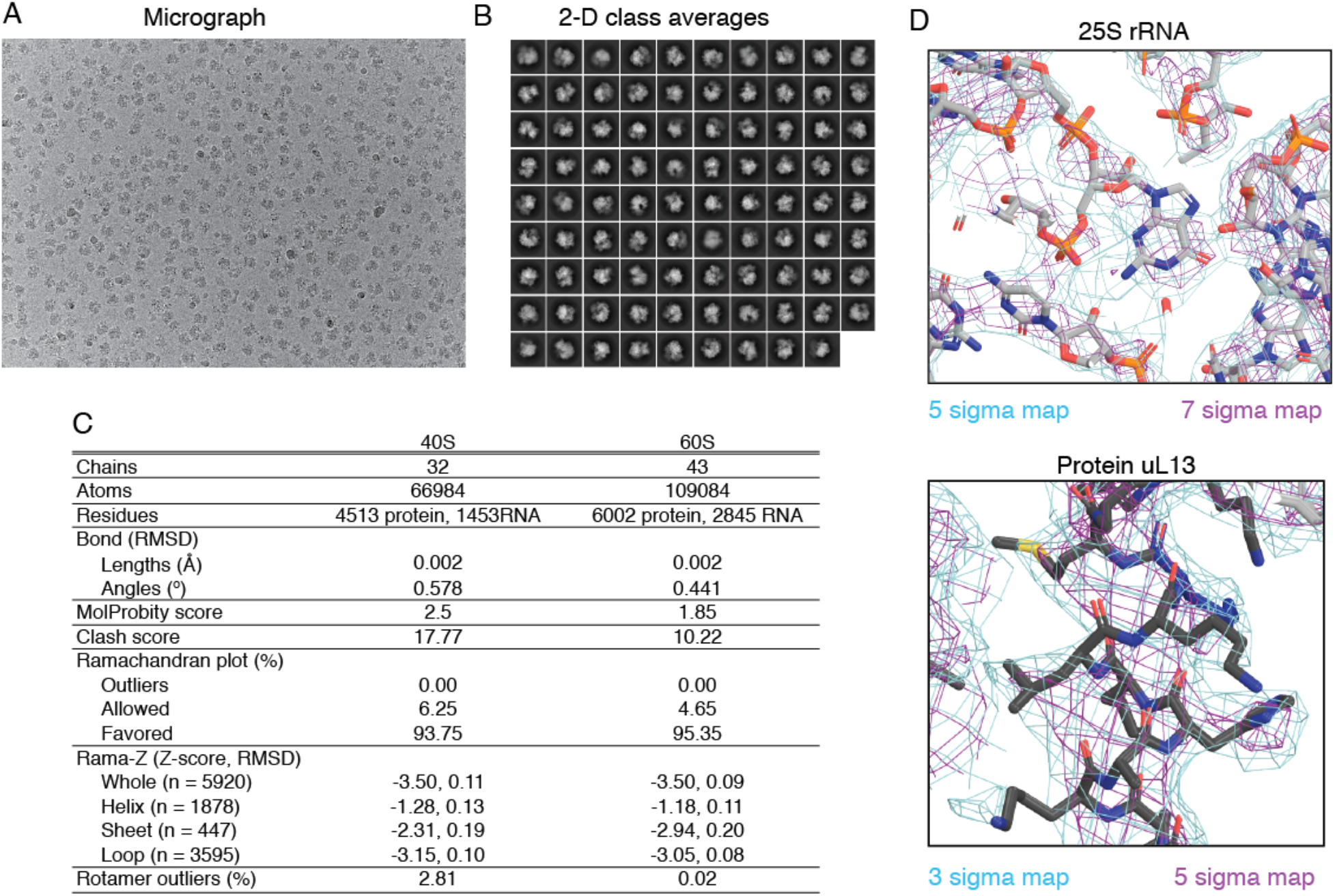
Details of cryo-EM. (A) Sample micrograph of the 80S ribosome from *G. lamblia*. Pixel size 1.39 Å. (B) 2D Class averages of the 80S ribosome from *G. lamblia*. (C) Refinement statistics of the structure of the 80S ribosome from *G. lamblia*. (D) Sample density around the 25S rRNA (left) and uL13 r-protein (right).

**Figure 1-Figure Supplement 2.**
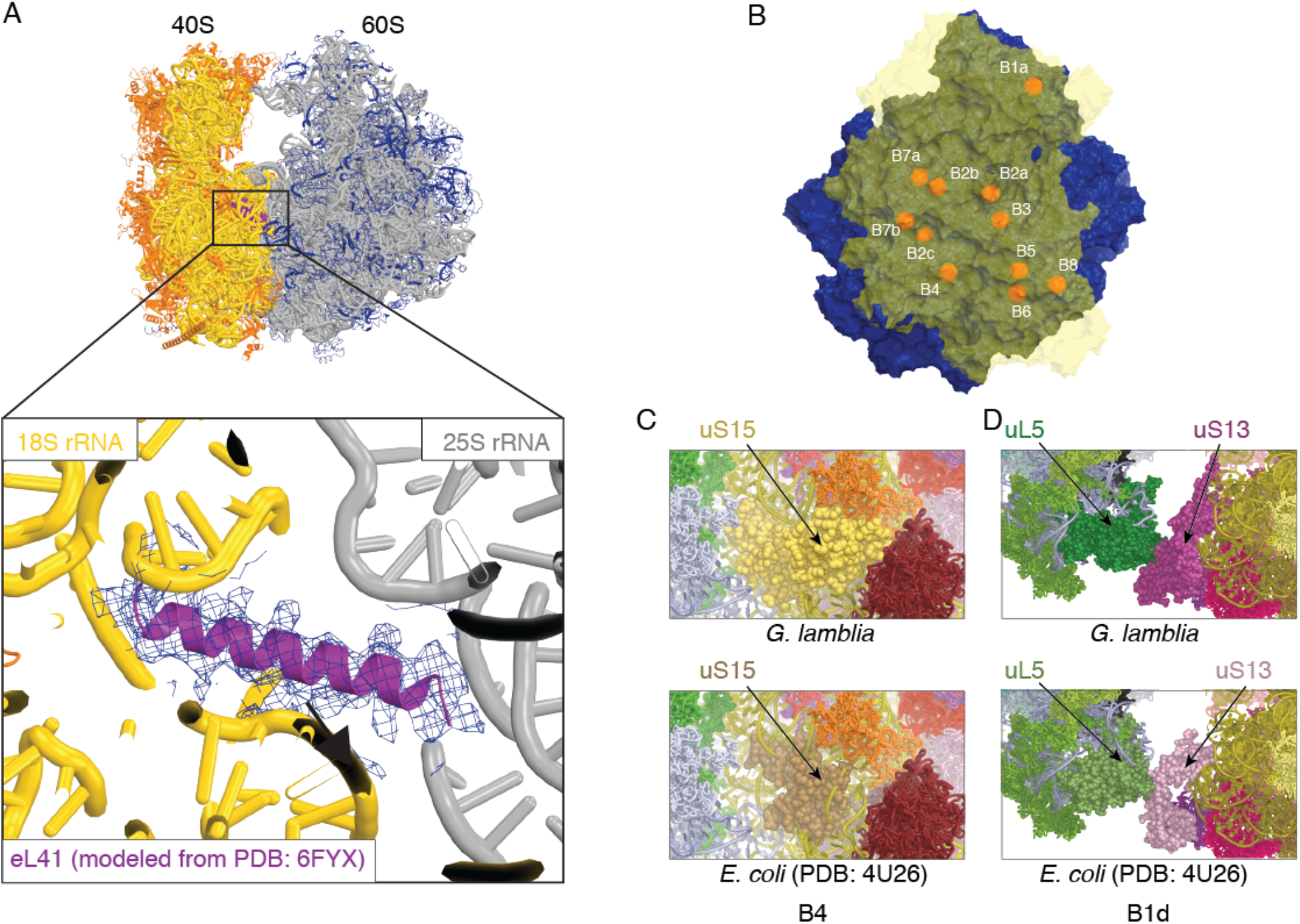
Intersubunit features of the *G. lamblia* 80S ribosome. (A) Density and structure of eL41 (purple) bound at the subunit interface. 40S subunit is in yellow and orange, 60S subunit is in gray and blue. eL41 from *S. cerevisiae* (PDB 6FYX) was used as there is no annotated sequence for *G. lamblia* eL41. (B) The *G. lamblia* 80S ribosome viewed from the solvent side of the 40S subunit, which is shown as a transparent yellow surface. Intersubunit bridges are indicated by orange spheres and labeled. The common set of intersubunit bridges B1a, B2a, B2b, B2c, B3, B4, B5, B6, B7a, B7b/c, and B8 are present in *G. lamblia*, although it lacks bridge B1b/c. In contrast, eukaryotic-specific bridges eB8 and eB11 are absent and eB12 is smaller, as expected, as these are associated with missing or shorter ESs. Interestingly, eukaryotic-specific bridge eB13, composed of protein, is present as is eB14, which in other eukaryotes is an interaction between eL41 and the 18S rRNA. (C) uS15 adopts a different conformation in *G. lamblia* (top, yellow) compared to *E. coli* (bottom, brown) to make bridge B4. D) uL5 (green) and uS13 (purple/pink) shown have an expanded interface in *G. lamblia* (top) compared to *E. coli* (bottom) to make bridge B1b.

**Figure 2–Figure Supplement 1.**
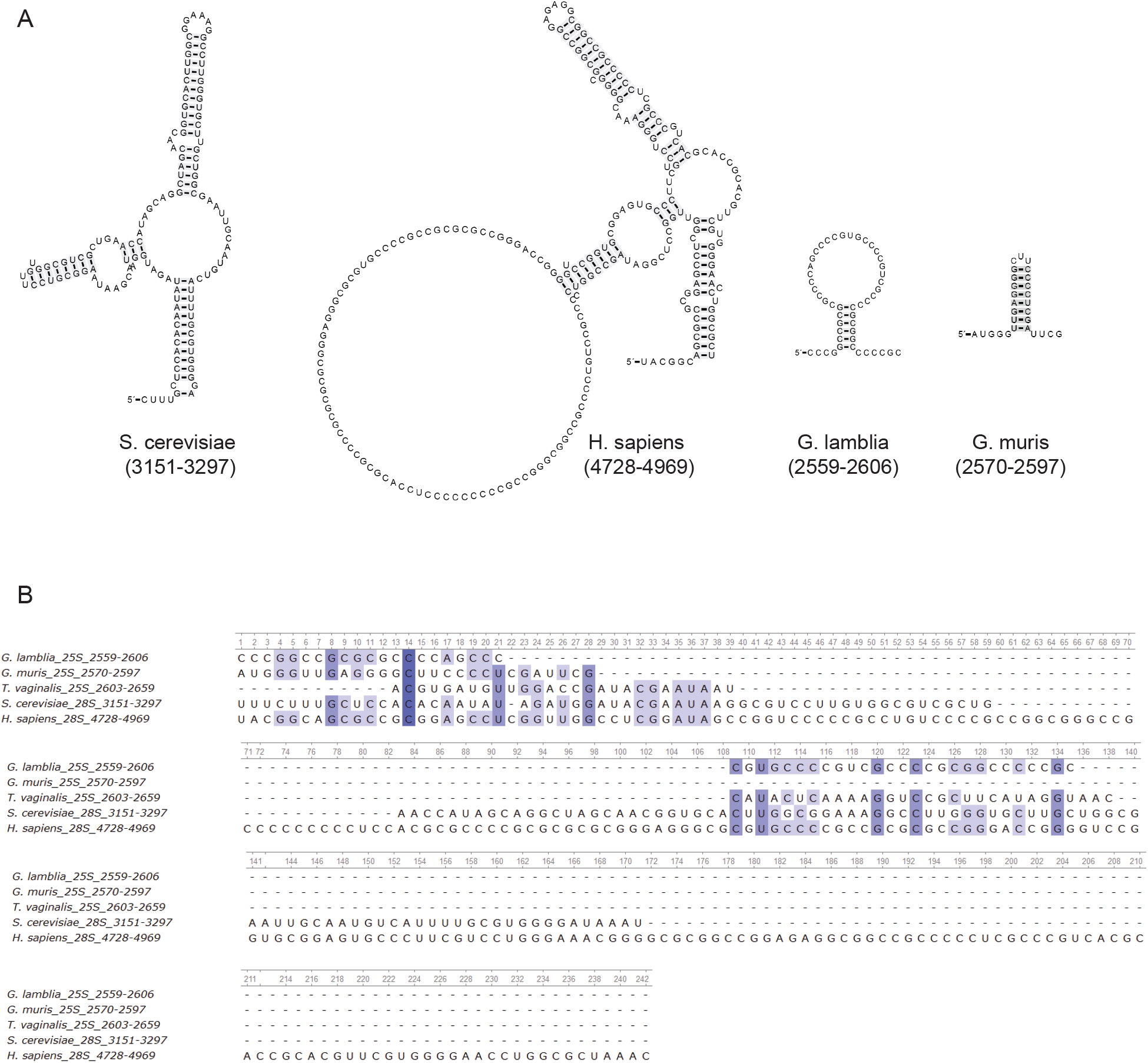
Secondary structure of divergent rRNA regions. (A) Secondary structure of the 25S/28S rRNA regions from Figure 2B. (B) Alignment of *G. lamblia, T. vaginalis, S. cerevisiae*, and *H. sapiens* 25S/28S rRNA regions from Figure 2B, using MUSCLE (Madeira et al., 2019).

**Figure 2–Figure Supplement 2.**
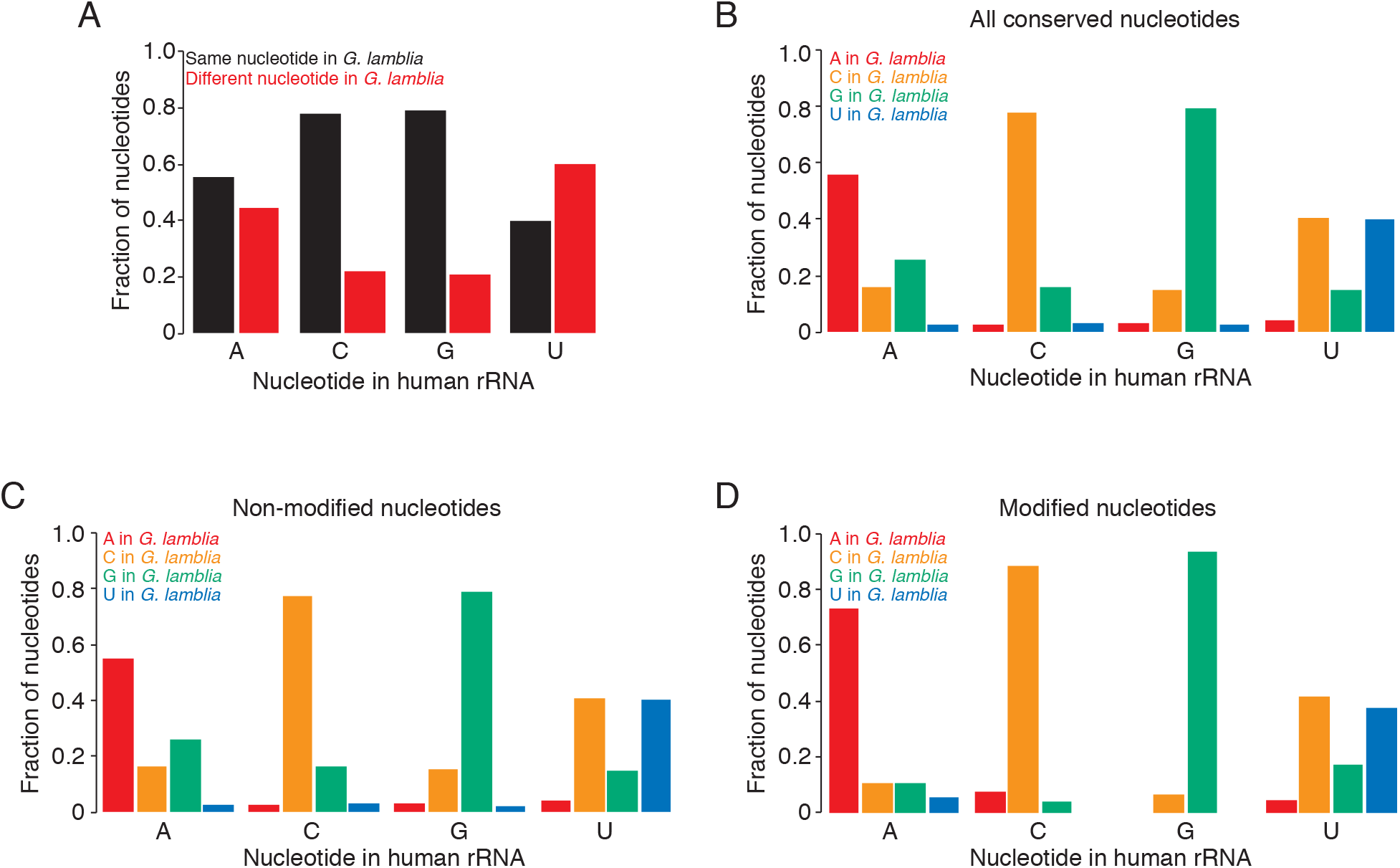
Analysis of nucleotide composition of G. lamblia rRNA. (A) Adenines and uridines in human rRNA are often different in *G. lamblia*. For rRNA nucleotides conserved between humans and *G. lamblia*, shown is the fraction of conserved nucleotides that are the same (black) or different (red) between the two. Nucleotides are grouped by their identity in humans. (B) Uridines are most often changed to cytidines in *G. lamblia* rRNA. For rRNA nucleotides conserved between humans and *G. lamblia*, shown is the identity of the nucleotide in *G. lamblia*, grouped by their identity in humans. (C) Non-modified uridines are often changed to cytidines in *G. lamblia* rRNA. As in B, but now only considering nucleotides non-modified in humans. (D) Modified uridines are often changed to cytidines in *G. lamblia* rRNA. As in B, but now only considering nucleotides modified in humans.

**Figure 3–Figure Supplement 1.**
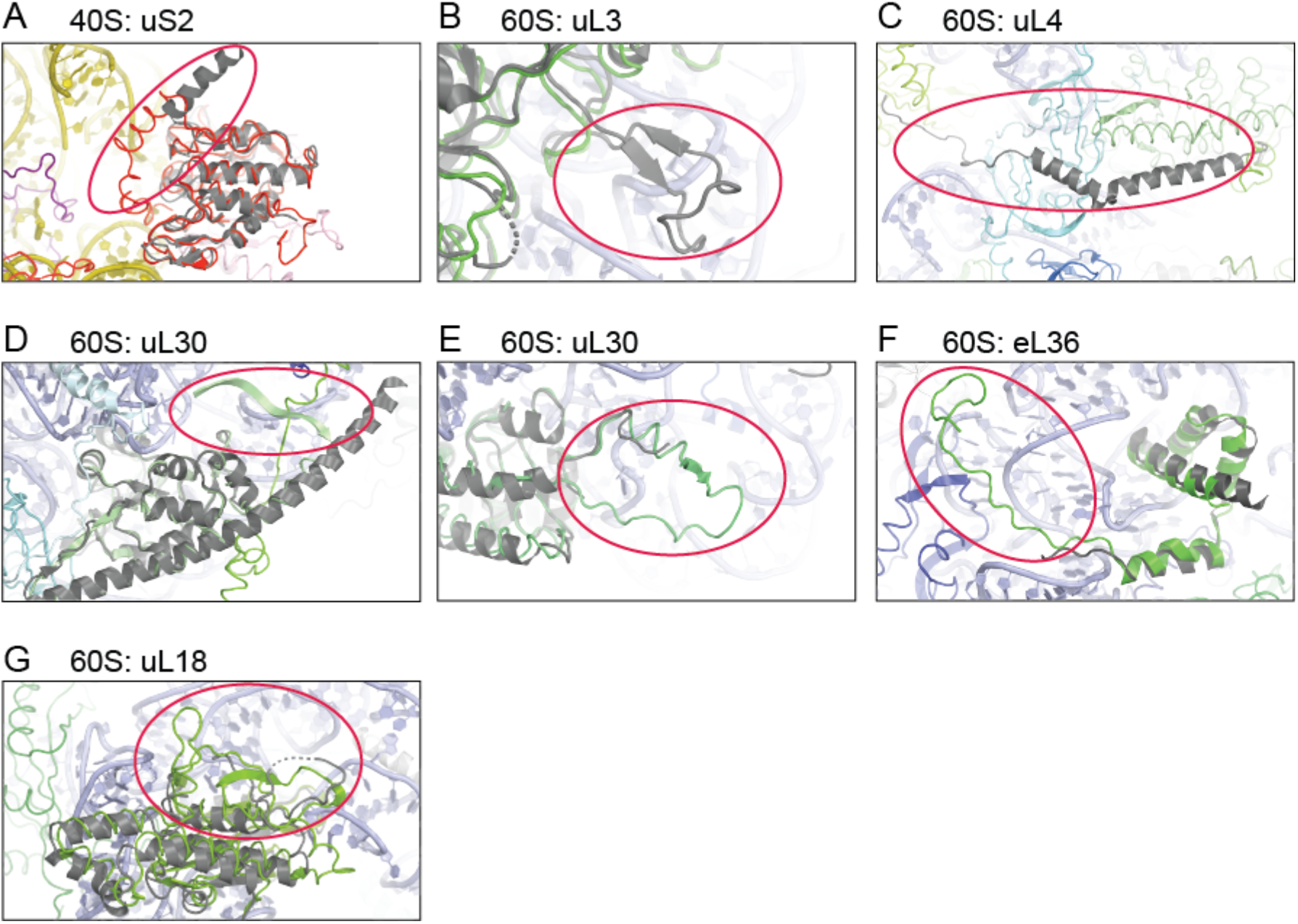
Changes in *G. lamblia* ribosome protein structures relative to *T. vaginalis*. In all panels, *G. lamblia* structures are colored and *T. vaginalis* are dark gray. (A) The C-terminal tail of uS2 on the 40S folds back into the body of the protein rather than into the cytoplasm. (B) uL3 is near the factor-binding stalk and has an internal loop deletion. C) uL4 has a large truncation of 31 amino acid in its C-terminus that wraps around the solvent side of the LSU. (D) uL30 is bound near the uL1 stalk and has an internal loop insertion and its N-terminus adopts a β-sheet rather than an α-helix fold; it still interacts with the 25S rRNA. (E) uL30 also has an internal loop expansion. (F) eL36 is found near the uL1 stalk and folds back to the core of the large subunit rather than being extended and interacting with ES5L, which is not present in *G. lamblia*. eL36 is found near the uL1 stalk and has a long C-terminal extension. (G) uL18 binds the solvent side of the LSU and has an internal loop insertion.

**Figure 4–Figure Supplement 1.**
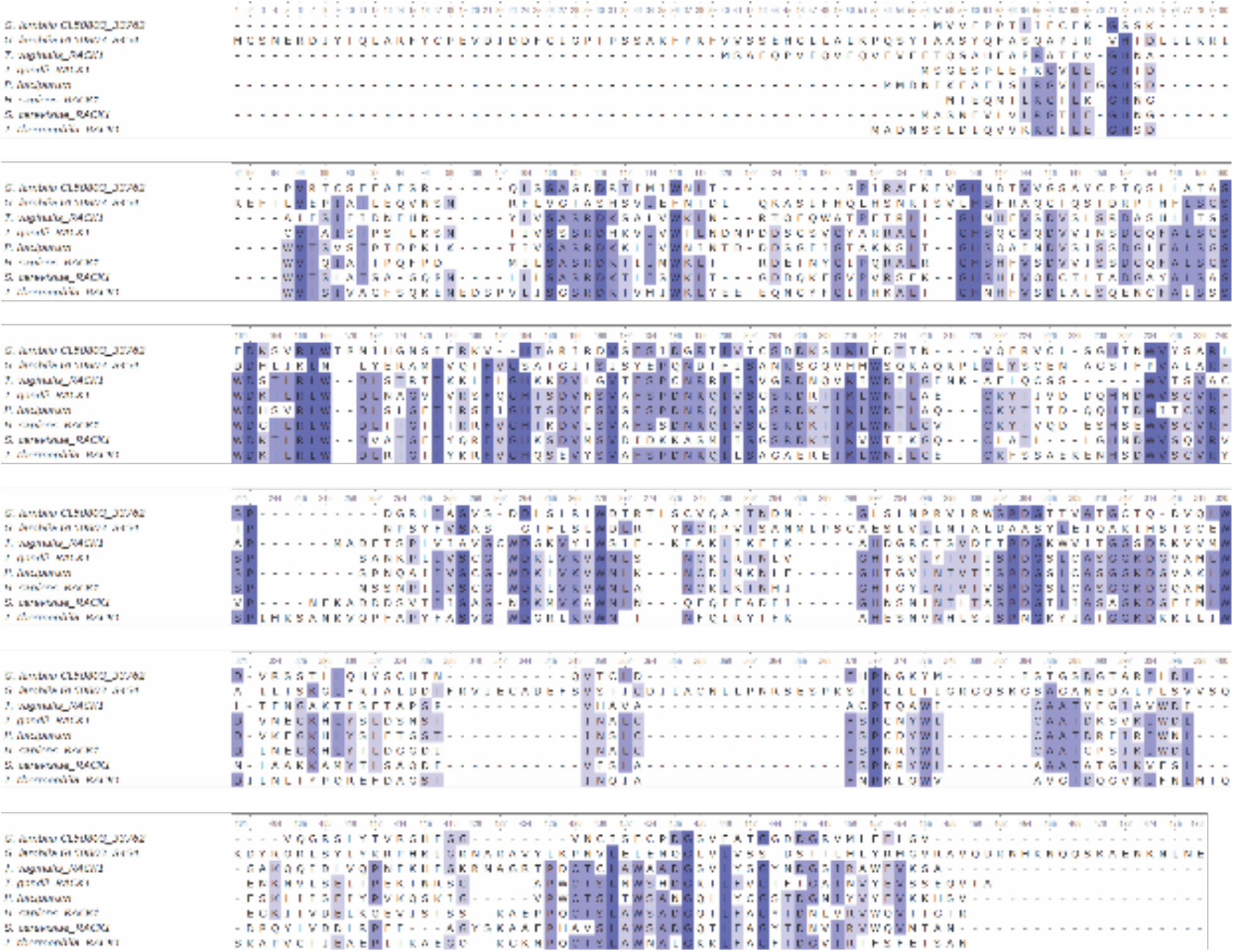
Alignment of RACK1 sequences. Sequences from *H. sapiens, S. cerevisiae*, *T. thermophila*, *T. vaginalis*, *G. lamblia*, *P. falciparum*, and *T. gondii* were aligned using MUSCLE (Madeira et al., 2019).

**Figure 4–Figure Supplement 2.**
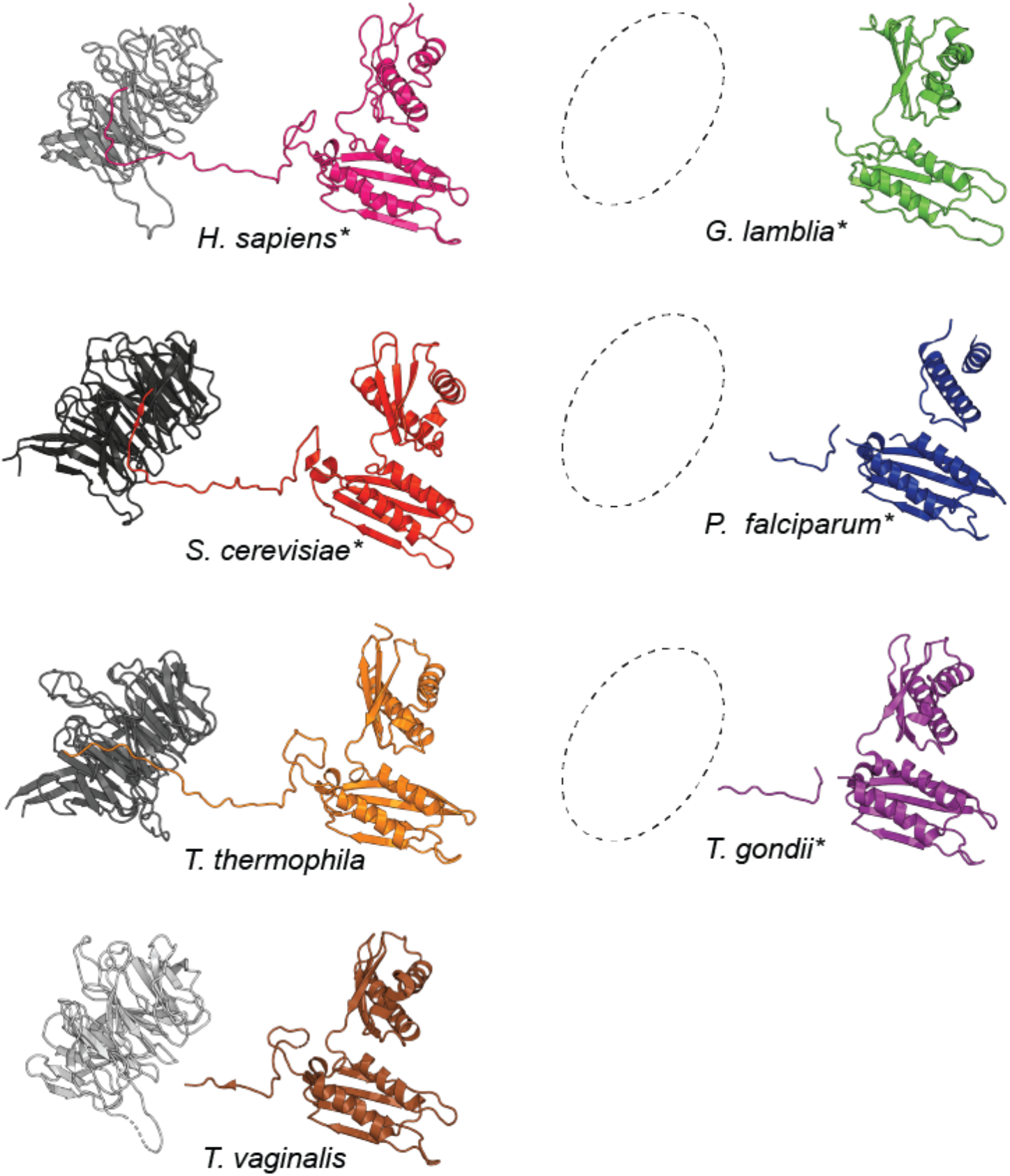
Differences in the presence of RACK1 and its interactions with uS3. The structures of RACK1 and uS3 bound to the ribosome for *H. sapiens*, *S. cerevisiae*, *T. thermophila, T. vaginalis, G. lamblia, P. falciparum*, and *T. gondii*. * indicates that there were amino acids disordered at the C-terminus and/or internal loop of uS3. Regions of disordered amino acids are indicated in Figure 4—figure supplement 3. Dashed open circles indicate that lack of RACK1 in the *G. lamblia*, *P. falciparum*, and *T. gondii* structures.

**Figure 4–Figure Supplement 3.**
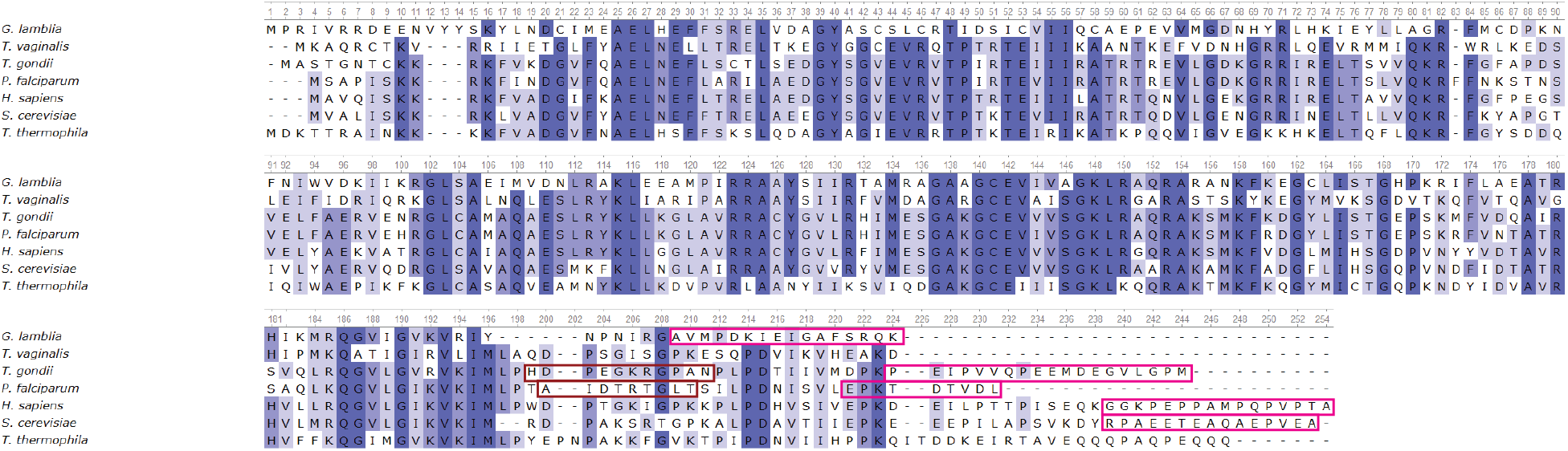
Alignment of uS3 sequences. Sequences from *H. sapiens, S. cerevisiae, T. thermophila, T. vaginalis, G. lamblia, P. falciparum*, and *T. gondii* were aligned using MUSCLE (Madeira et al., 2019). Amino acids disordered at the C-terminus of uS3 are indicated with magenta boxes and amino acids indicated with dark red are disordered internal loops.

**Figure 4–Figure Supplement 4. Gene identification number of potential NGD/RQC orthologs**. Table contains gene identification numbers of most likely orthologs (base on sequence homology) for No-Go Decay factors in various protist species.

Supplied as a separate spreadsheet file.

